# Adhesion-free cell migration by topography-based force transduction

**DOI:** 10.1101/793919

**Authors:** Anne Reversat, Jack Merrin, Robert Hauschild, Ingrid de Vries, Matthieu Piel, Andrew Callan-Jones, Raphael Voituriez, Michael Sixt

## Abstract

Eukaryotic cells migrate by coupling the intracellular force of the actin cytoskeleton to the environment. While force-coupling is usually mediated by transmembrane adhesion receptors, especially these of the integrin family, amoeboid cells like leukocytes can migrate extremely fast despite very low adhesive forces^1^. We show that leukocytes cannot only migrate under low adhesion but indeed can transduce forces in the complete absence of transmembrane force coupling. When confined within three-dimensional environments, they use the topographic features of the substrate to propel themselves. Here, the retrograde flow of the actin cytoskeleton follows the texture of the substrate, creating shear forces sufficient to drive deformations towards the back of the cell. Notably, adhesion dependent and adhesion independent migration are not exclusive but rather variants of the same principle of coupling retrograde actin flow to the environment and thus can potentially operate simultaneously. As adhesion free migration is independent of the chemical composition of the environment it renders cells completely autonomous in their locomotive behavior.

## Main Text

Mechanistic understanding of cell motility started with Abercrombies demonstration that particles placed on the dorsal surface of a cell undergo retrograde transport ^2^. The authors suggested that “cellular material” is continuously added to the front of the leading edge and generates a rearward flow, which couples to the substrate and thereby generates friction to drag the cell forward. Later studies showed that the added material is the growing polymer of F-actin and that F-actin retrograde flow couples via transmembrane adhesion receptors, mainly of the integrin family. Although this adhesion-dependent principle of motility turned out to be universal on two dimensional (2D) surfaces, cells confined within a 3D context often migrate in the absence of integrin mediated adhesion ^3,4^. This applies especially to the amoeboid (shape-changing) class of cells like leukocytes ^5^. How these cells generate friction with their environment remains unknown, and alternative membrane spanning molecules have been suggested ^6,7^. Interestingly, in his seminal study, Abercrombie contemplated a second, qualitatively different, scenario: backwards moving waves of deformation might “passively” entangle with the particles (or a substrate) without adhering and propel the cell in a manner analogous to swimming ^2^. And indeed, some amoeba were shown to “swim” towards chemoattractants when buoyant in viscous media ^8–10^.

To address the question how adhesion-independent locomotion might function we employed a murine T cell line, which migrates vigorously in 3D environments like *in vitro* assembled collagen gels (Fig. 1c and Extended Data Fig. 1e and supplementary video 1). In order to eliminate integrin-based force transduction, we deleted talin 1, a cytosolic adaptor protein essential for integrin functionality (Extended Data Fig. 1b and ref). Talin-deficient cells (“talin KO”) were completely unable to adhere to and migrate on 2D surfaces (Fig. 1a, b and Extended Data Fig. 1c-1d). However, once incorporated into 3D collagen gels, their migratory characteristics were indistinguishable from wild type control cells (Fig. 1c and Extended Data Fig. 1e and supplementary video 1 and ^5^). In the following, we re-engineered artificial environments to dissect the geometrical transition between 2D surfaces and complex fibrillar environments, while keeping the chemical composition of the surface constant. We first designed a microfluidic setup to confine leukocytes between two parallel surfaces separated by a 5 µm space (“2,5D”, ^11^ and Fig. 1d). While this setup supported efficient migration of the wild type T cells (^3^, Fig. 1d, Extended Data Fig. 1f and supplementary video 2), talin KO cells were hardly migratory (^3^, Fig. 1d, Extended Data Fig. 1f and supplementary video 2), although they showed morphological front back polarization (supplementary video 2). When T cells expressing the actin reporter Lifeact-GFP were imaged with total internal reflection (TIRF) microscopy, control cells showed substantial retrograde flow in the lamellipodial region, while below the cell body, actin was static in relation to the substrate (Fig. 1e, f and g and supplementary video 3). Talin KO cells showed similar actin dynamics in the cell frame of reference, but as they did not move, all actin flowed backwards in relation to the substrate (Fig. 1e, f and g and supplementary video 3). This was true for different degrees of confinement and even in narrow (3 µm) spaces, when cells formed large dynamic blebs (as shown in ^3^) talin KO cells showed no forward locomotion (Fig. 2f, g, Extended Data Fig. 1h and supplementary video 4). These findings demonstrate that, when confined between two surfaces, the cells transduce forces in an integrin-dependent manner. Deletion of talin 1 completely decoupled cytoskeleton and substrate, but left the cytoskeletal dynamics of the cell unaffected, essentially causing it to run on the spot. This suggested that sheer confinement does not create sufficient friction to drive locomotion.

**Figure 1.**
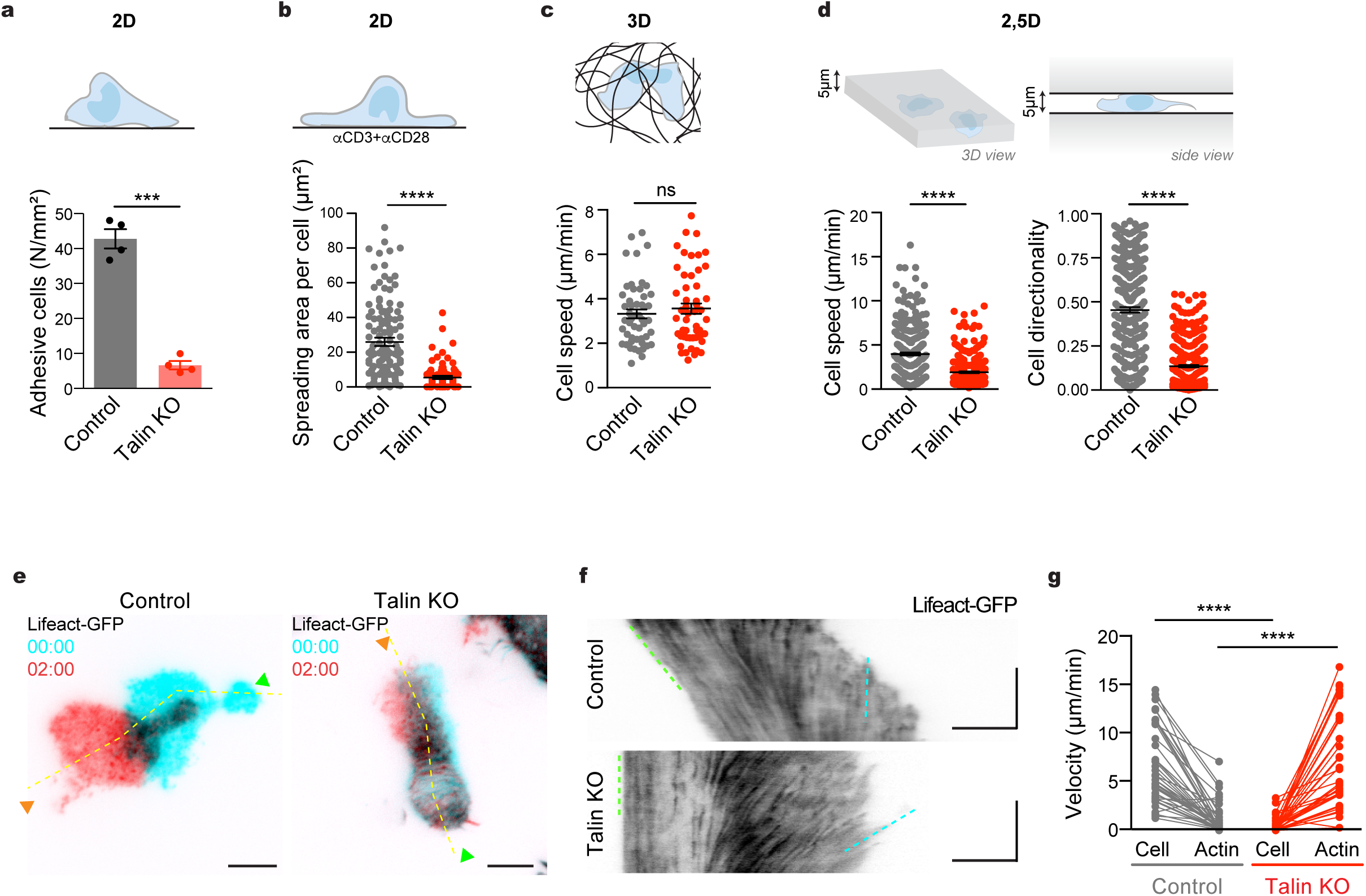
Adhesive and migratory properties in 2D, 3D and 2,5D environments of control and talin KO leukocytes. **a**, Upper panel: scheme of T cells on a 2D dish. Bottom panel: quantification of the number of T cells adhering to the cell culture dish; quantification from 4 independent experiments shows mean ± SEM (*** p=0.00076; paired *t*-test). **b**, Upper panel: schematic representation of the 2D spreading assay onto anti-CD3 and anti-CD28 coated glass. Bottom panel: quantification of the spreading area (µm^2^) observed in TIRF in control (n=139) and talin KO (n=81) T cells from 2 independent experiments (mean ± SEM is shown in black; **** p<0.0001; Mann-Whitney *U* test). (**c**) Upper panel: scheme of a collagen assay for cell migration in 3D. Bottom panel shows the individual speed of control (n=51) and talin KO (n=52) T cells in 3D collagen gel observed in 3 independent experiments (mean ± SEM in black; p=0,628; Mann-Whitney *U* test). **d**, Upper panel: scheme of the 2,5D cell confinement used in (d-g). Bottom panel: the quantification of migration speed (c) and directionality (d) of control (n=290) and talin KO (n=284) T cells in 5 µm confiner observed in 4 independent experiments (mean ± SEM is shown in black; p<0.0001; Mann-Whitney *U* test). **e**, Snapshots of TIRF microscopy of control (left panel) and talin KO cells (right) expressing Lifeact-GFP reporter under 5 µm height confinement. Green arrows show the contractile uropod of the T-cell back, blue arrow the cell front. TIRF is shown in cyan at t=0min, and red at t=2min, and the yellow dotted line is used for kymograph analysis in f. Scale bar, 5 µm. Time in min:sec. **f**, Kymographs along back-front polarity of control and talin KO T cell under 5 µm confinement. Cell velocity is indicated with a green dotted line, and actin retrograde flow in cyan. Horizontal scale bar, 5 µm, vertical 1min. **g**, Measurement of T cell and actin retrograde flow velocities at a single-cell level, in control (n=42) and talin KO (n=36) (3 independent experiments; **** p<0.0001; Mann-Whitney *U* test).

**Figure 2.**
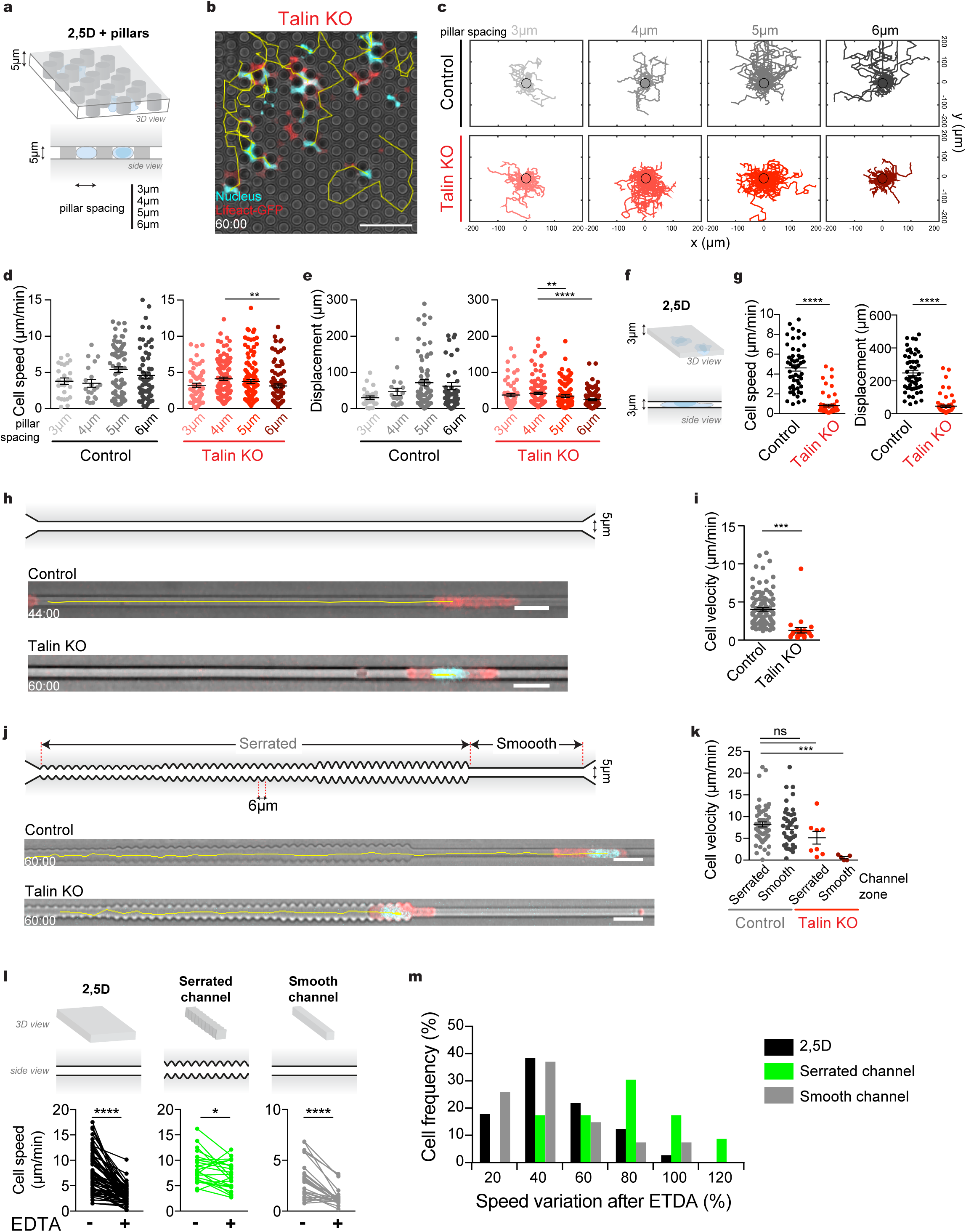
Migratory properties of leukocytes in topography-bearing confinement. **a**, Scheme of the pillar maze used in b-i. 10 µm diameter pillars separated by either 3, 4, 5 or 6 µm are added to the 2,5D cell confinement. (**b**) Snapshots of a video microscopy at t=60min of talin KO T cells in the 4 µm-spaced pillar maze device. Cyan shows the nucleus (Hoechst), red the Lifeact-GFP reporter, and gray the brightfield. Scale bar, 50 µm. Time in min:sec. **c**, Control and talin KO T cells trajectories in the maze device during 1 hour. **d**, Quantification of cell speed (mean ± SEM; ** p=0.0021 for talin KO otherwise not significant; Kruskal–Wallis followed by a post hoc Dunn test). **e**, Cell displacement in the maze (mean ± SEM; ** p=0.0089; **** p<0.0001 otherwise not significant; Kruskal–Wallis followed by a post hoc Dunn test). n=25, 20, 75 and 61 for control (3 independent experiments) and n=64,115,124 and 94 for talin KO T cells in the respective 3, 4-, 5-, and 6 µm pillar spacing zone (6 independent experiments). **f**, Scheme of the 3 µm high 2,5D cell confinement. **g**, Quantification of cell speed and displacement of control and talin KO in the 3 µm confinement from 4 independents experiments, n=52 for control, n=60 for talin KO (mean ± SEM; **** p<0.0001; Mann-Whitney *U* test). **h**, Scheme of the 5×5µm smooth microchannel (upper panel) and snapshot at t=44min of Lifeact-GFP Control (middle panel) and at t=60min Lifeact-GFP talin KO (lower panel) T cells. Tracks are display in yellow, Scale bar 20 µm. **i**, Quantification of cell velocity in smooth channels from 4 independents experiments, n=99 for control, n=24 for talin KO (mean ± SEM ; **** p<0.0001; Mann-Whitney *U* test). **j**, Scheme of the serrated microchannel (upper panel). Serrated topography with a periodicity of 6 µm were introduced into a 5×5 µm microchannel. Snapshot at t=60min of Lifeact-GFP Control (midlle panel) and talin KO (lower panel) T cells in microchannels baring 6 µm-spaced geometries. **k**, Single cell velocities (µm/min) of control and talin KO T cell, migrating in serrated microchannels. The cell velocity was measured separately in the serration zone or the smooth zone; n=59 (control serrated), 25 (control smooth), 8 (talin KO serrated) and 5 (talin KO smooth) (mean ± SEM is shown; *** p=0.0009; Kruskal–Wallis followed by a post hoc Dunn test). **l**, Control T cells migrate into a device containing 2,5D confinement, smooth and serrated microchannels as shown in upper panels. After 60 minutes 10mM EDTA is added and single-cell migration speed is observed before (-) and after (+) EDTA treatment. 3 independent experiments, (n=73 for 2,5D with **** p<0,0001; serrated channel n=23 and * p=0,0167; smooth channel n=27 **** p<0,0001; Wilcoxon matched-pairs signed rank test). **m**, Single-cell variation of migration speed after vs before EDTA treatment.

We reasoned that the major difference between confinement within a fibrillar matrix and between two smooth surfaces is geometrical complexity. Hence we introduced arrays of 10 µm diameter pillars intersecting the two surfaces. Pillar spacing ranged between 3 and 6 µm (Fig. 2a). Notably, talin KO cells were unable to actively enter the area of confinement, while control cells could do so. This is consistent with our finding that integrins are essential for invasion of 3D matrices but not for locomotion within them (data not shown). We thus used a valve-containing pressure-controlled microfluidic set-up to gently push cells into the confined area (Extended Data Fig. 2a and materials and methods). Remarkably, once the talin KO T cells were positioned between the pillars, their migratory capacity was restored (Fig. 2b, c, d, e, Extended Data Fig. 2b and supplementary video 5). Quantification of single cell velocities revealed that compared to 3-4 µm, within 6 µm spaced pillars the performance of talin KO cells dropped significantly, suggesting that an optimal pillar spacing enables fast migration in the absence of integrin mediated force coupling (movies S5).

The sharp transition between no locomotion in 2,5D confinement and restored locomotion in the pillar arrays provides evidence that geometrical rather than chemical features suffice to generate propulsion forces between cells and substrate ^12^. However, the pillar setup did not exclude that the increased surface contact area might provide more “unspecific friction” by non-integrin transmembrane receptors. To address this caveat, we switched to a more reductionist setup and designed microchannels where cells are in full contact with the channel walls (Fig. 2h). Within smooth-walled 5×5 µm microchannels (Fig. Extended Data Fig. 2a) control cells migrated persistently, while talin KO cells were immobile (Fig. 2h, i and supplementary video 6). We next introduced geometrical irregularities as a serrated pattern with a 6 µm period (Fig. 2j, k, Extended Data Fig. 2c and supplementary video 7) and found that migration of talin KO cells was restored. In microchannels containing both smooth and serrate sectors, wildtype cells continuously transited between the sectors. Upon cation depletion by EDTA, which inactivates integrins and several other cell surface adhesion molecules, cells within smooth areas slowed or stalled, while cells within textured sectors kept migrating until they encountered a smooth area. (Fig. 2l and 2m and supplementary video 8).

As a framework to understand topography based amoeboid locomotion, we propose a minimal model (Fig. 3 and supplementary text) where actin is described as a viscous gel travelling from the leading to the trailing edge of the cell in absence of any tangential friction force with the substrate. In serrated but not in smooth channels this flow generates local shear stresses, and thus a pressure gradient in the actin gel, driving locomotion by inducing non vanishing normal forces onto the substrate. Here, the actin flow generated in the cell can couple to the environment when its topographic features are smaller than the flow scale. To test these predictions and generate varying length scales, we varied the period of serrated topography from 6 µm to 12 µm to 24 µm. Serrated sectors were followed by smooth areas as an internal control (Fig. 4a). Wildtype cells effectively traversed all channel designs, while talin KO cells migrated in 6 and 12 µm period patterns but rarely in the 24 µm patterns (Fig 4b, c and d), where the spacing exceeded the average length of cortical actin flow in a cell (Extended Data Fig. 2f-g). In channels where the period successively increased from 6, to 12, 24 µm, continuous single cell observation showed that cells gradually slowed, showed saltatory movement in the 24 µm stretches and (Fig. 4c, d and supplementary video 9) were unable to enter the smooth channel, in qualitative agreement with model predictions. We next used confocal and TIRF imaging of Lifeact-GFP-expressing talin KO cells in topographic microchannels. Kymograph analysis revealed an increase of actin retrograde flow whenever the cells slowed between the topographical features (Fig. 4e, f and g and supplementary video 10), suggesting that, in analogy to the transmembrane clutch, topographical features can serve to couple retrograde actin flow to a substrate.

**Figure 3.**
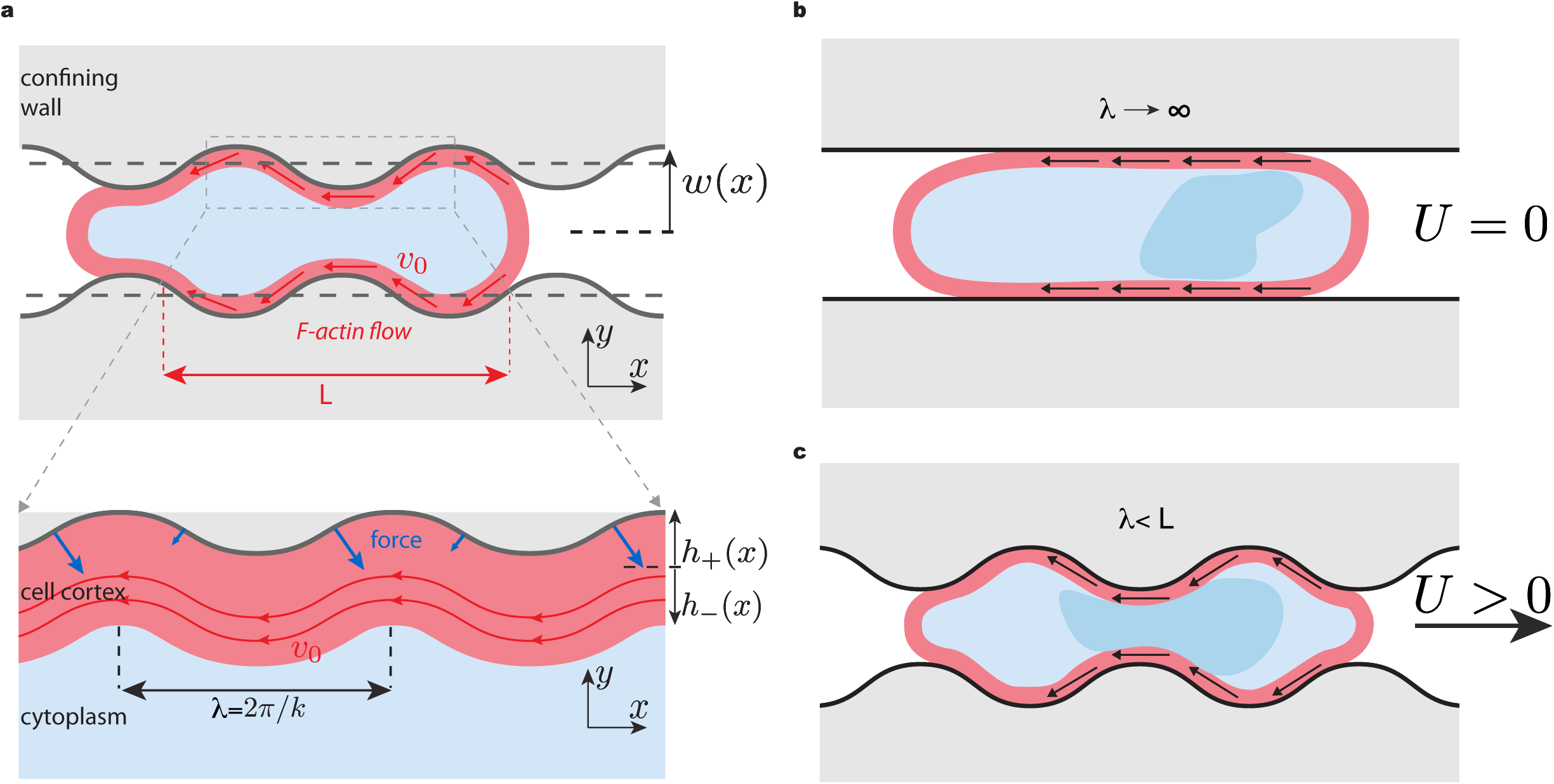
Physical model of adhesion-free cell migration in a complex environment. **a**, Insight into the model: the channel has a varying wavy cross-section w(x) and period λ=2π/k. The cell contains cortical F-actin that presents a retrograde flow of length L and average speed *v*0, and is modelled as a viscous fluid layer of thickness h(x). **b**, Scheme of a modelled cell in a smooth channel with no adhesive-based coupling of the actin cytoskeleton with the environment. The cell is polarized and has a high retrograde actin flow but cannot move forward. **c**, Scheme of the modelled non-adhesive cell in a serrated channel. Cell propulsion is created by the shear stress in the actin cortex, which is caused by the bending of flow lines by the environmental topography. This induces a pressure gradient in the actin gel and thus non vanishing normal forces onto the substrate.

**Fig. 4.**
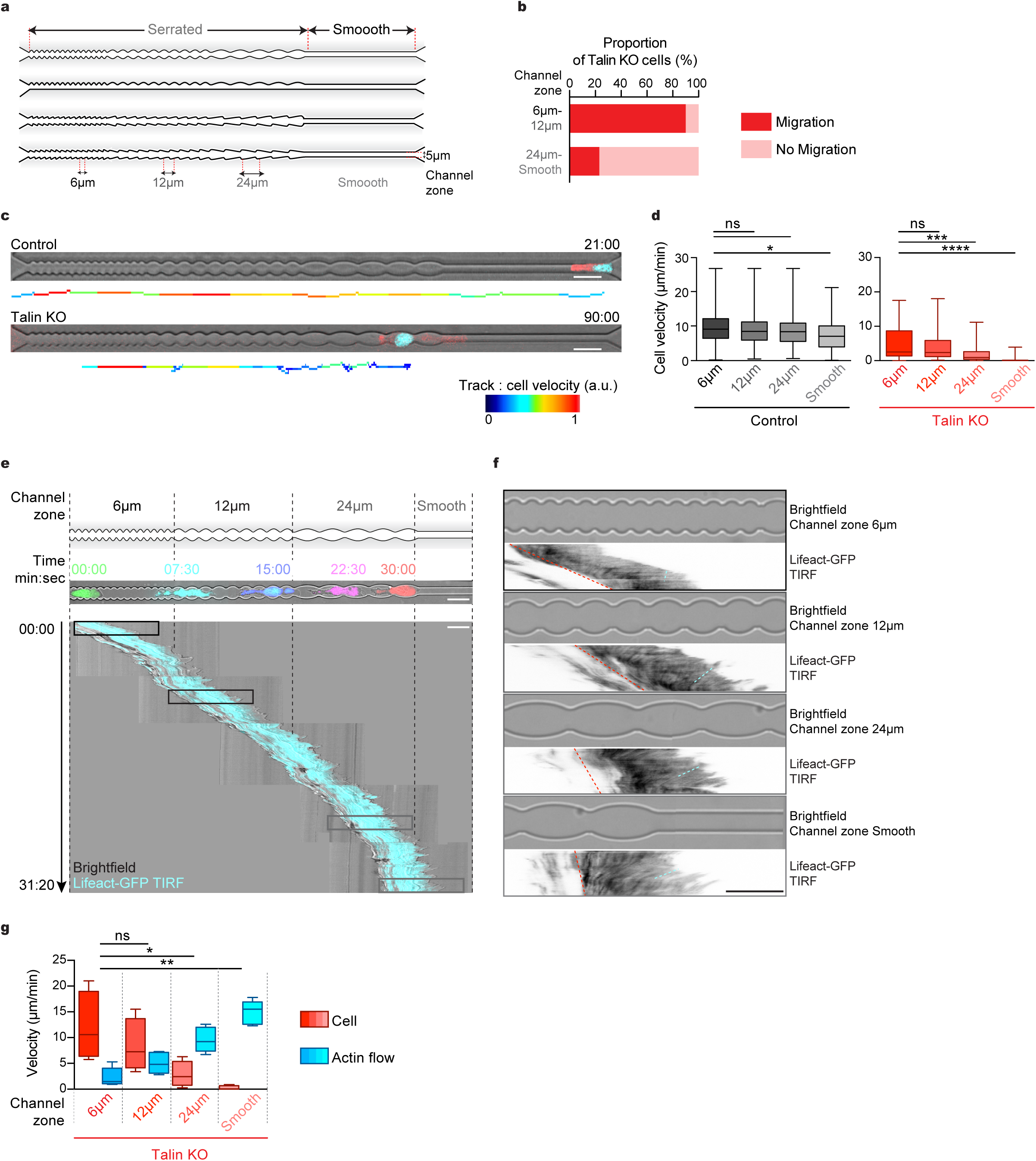
Dependency of talin KO leukocytes locomotion with the complexity of the geometry of the environment. **a**, Scheme of the microchannels designs with decreasing complexity used in b-g. Briefly, serrations with periodicity of 6, 12 and 24 µm and were put consecutives zones of a microchannel (5 µm wide), ending by a smooth zone. Different geometries were created, like the symmetric bulb and oriented arrows (cf figure S3). **b**, Migration capacity of talin KO cell in the 6 or 12 µm zone (n=41) and in the 24 and smooth zone(n=22). Fisher test; **** p<0,0001. **c**, Video microscopy snapshots of Control (upper panel, t=21min) and talin KO (lower panel, t=90min) T cells in microchannels baring periodic geometries. Lifeact is shown in red, Hoechst in cyan. Scale bar 20 µm. Below the snapshots tracks are color-coded for single cell velocity (In red maximum velocity of control 28,3 µm/min; talin KO 34,1 µm/min). **d**, Single-cell migration analysis into the different periodicity geometries. Left panel shows cell velocities for control (gray, n=88, 4 independent experiments) and talin KO T cells (red, n=79, 8 independent experiments) into the different channel zones (Control *p=0.0234; KO ** p=0.0013; **** p<0.0001 otherwise not significant; Kruskal–Wallis followed by a post hoc Dunn test). **e**, TIRF microscopy of migrating talin KO cells from the 6 µm zone to the entrance of the smooth zone. The cells express Lifeact-GFP reporter. Left panel, from top to bottom: scheme of the microchannel, color-coded snapshots every 7,5 min and kymograph of a cell migrating along the channel (cyan: TIRF, gray: brightfield). Scale bar 20 µm, time display in min:sec. **f**, Brightfield of the different channel zones and corresponding kymograph of F-actin TIRF of a talin KO cell. x scale bars 20 µm, y of kymograph corresponds to 50s. **g**, Quantification of talin KO cell velocities and actin retrograde flow velocities observed by TIRF microscopy and measured at a single cell-level with kymograph analysis (n=7; 3 cells in all zone, 2 cells in 6-12-24 µm zone, and 2 cells in smooth zone ; * p=0.0423 ; ** p=0.0036 Kruskal–Wallis followed by a post hoc Dunn test). Time in min:sec.

Altogether, our findings suggest that cells can transduce forces by coupling the retrograde flow of actin to a geometrically irregular environment, and this can happen in the complete absence of any transmembrane receptors linking the cytoskeleton to the substrate. Most likely, adhesion dependent and adhesion independent migration are not exclusive modes but two variants of the same fundamental principle of coupling actin flow that can operate simultaneously. We demonstrated this principle by controlling the topography of the environment, thereby imposing shape changes extrinsically. Notably, amoeboid cells have the intrinsic capacity to produce rearward propagating shape waves ^9,10,13^, which, once entangled with a substrate, can propel the cell, as suggested by Abercrombie.

## Supporting information

Supplementary Model

## Acknowledgements

We thank Alexander Leithner, Joerg Renkawitz for discussion and critical reading of the manuscript, Jan Schwarz and Matthias Mehling for the establishment of the microfluidic set-up, the Bioimaging Facility of IST Austria for excellent support, as well as the Life Science Facility and the Miba Machine Shop of IST Austria. A.R. thanks Feyza Nur Arslan, Laura Elisabeth Burnett and Lanxin Li for their work during their rotation in the ISTScholar PhD program. This work was supported by the European Research Council (grant ERC GA 281556) to M.S. and a START award from the Austrian Science Foundation (FWF) to M.S.

## Author Contributions

AR and MS conceived the experiments and wrote the manuscript. AR designed, performed, and analyzed experiments, with the help of IDV. AR and JM designed the microfluidic devices, and JM performed the photolithography. RH wrote image analysis scripts. ACJ, RV, JM, RH, MS and AR discussed about the physical model and ACJ, RV wrote the physical model.

## Supplementary Materials

Supplementary Text: Model

Materials and Methods

Supplementary Table S1

Extended Data Fig. 1 and 2

Captions for Extended Data Fig. 1 and 2

Supplementary videos 1 to 10

Captions for supplementary videos 1 to 10

