## Supplementary Model for "Adhesion-free cell migration by topography-based force transduction"

### Supplementary Text: Model of Adhesion-free cell migration by topography-based force transduction

#### I. INTRODUCTION

We describe here a simple model of cell migration, which shows that a confined cell can be efficiently motile in absence of any adhesion or friction-like interaction (specific or non specific) with its environment, provided that the environmental geometry is sufficiently irregular. The model is general and does not rely on a specific cell type; its aim is to identify the minimal ingredients required for adhesion-free cell migration. As we show below, the main ingredient is a polarised flow of the actin cytoskeleton, which is a very conserved feature of motile cells and was shown in various contexts and cell types to be responsible for cell motion[3,15–17]. We do not describe here the molecular mechanisms generating the actin flow, such as actin polymerisation/depolymerisation and/or myosin II induced contractility. Those are assumed to sustain a retrograde flow of average speed $v_0$ (note that here flows are defined in the lab reference frame) that extends along the cell polarity axis over a typical length $L$, which can be smaller than the full cell size. At the time scales of experiments (larger than minutes) viscoelastic effects are negligible and the actin cytoskeleton can be modelled as a fluid of viscosity[17], $\eta$.

Let us first recall that in absence of specific adhesion or friction, the only forces that the environment can exert on a cell are normal to the local cell boundary. This shows that the external forces experienced by cells confined between parallel planes or in channels with flat walls have necessarily a vanishing component along the cell polarity axis. Motion is therefore impossible in these cases, in agreement with observations (see main text). In irregular geometries, normal forces have a *locally* non vanishing component along the cell polarity axis. Whereas for a static fluid such normal forces are homogeneous and sum up to zero, we show here on general grounds that in the presence of a retrograde actin flow, they are inhomogeneous and sum up *globally* to a non vanishing propulsion force, which in turn enables migration. Qualitatively, an irregular geometry bends the actin flow lines and induces shear forces in the actin cytoskeleton. To maintain the actin flow and balance shear forces, a pressure gradient along the cell polarity axis is therefore required in irregular geometries, and locally induces non homogeneous normal forces : forward facing boundaries are subject to larger forces than backward facing boundaries, leading to an overall non vanishing forward force (see Fig. 3). Importantly, this locomotion mechanism does not require any large scale asymmetry of the environment (or ratchet like effect); the direction of motion is imposed solely by the cell polarity, or equivalently in the model by the direction of the actin flow, which we assume is set independently of the environment. In particular, while we consider below sinusoidal shapes of the confining environment for the sake of simplicity, the mechanism that we describe is applicable to any irregular geometry.

#### II. MODEL AND SUMMARY OF THE RESULTS

To justify more quantitatively this generic mechanism we take the example of a channel with modulated width used in experiments, as in Fig. 3. The channel cross-section varies with the longitudinal coordinate $x$ and can be written mathematically as $w(x) = w_0 + w_1 \sin(kx)$; here $\lambda = 2\pi/k$ is the wavelength of the wall deformation and the local depth of the channel in contact with the cell, $z_0$, is assumed constant. We focus on the cell subregion of length $L$ where an actin retrograde flow of average speed $v_0$ in the lab frame (here $v_0 < 0$) along the $x$ axis takes place. Actin is assumed to be localised in a cortical layer of thickness $\delta$ that covers the walls of the channel; following the above discussion a non trivial contribution to cell motion can come only from the cortex layers facing the wavy walls.

For simplicity, in the derivation below we assume that $L$ is a multiple of the spatial period, i.e., $L = n\lambda$, so that we can consider a periodic geometry; the generic case where $L/\lambda$ is arbitrary is discussed below. We therefore consider a slab of cortex delimited by $0 < x < L$ and $w(x) - \delta < y < w(x)$, and we neglect the dependence on the normal coordinate $z$. For convenience we introduce local coordinates and take $y = w_0 - \delta/2$ as a new reference axis; the cortex layer of interest then extends from $y = -h_-(x) = -h_0 + (1 - \epsilon \sin(kx))$ to $y = h_+(x) = h_0(1 + \epsilon \sin(kx))$, where $h_0 = \delta/2$ and $h_0\epsilon = w_1$.

##### A. Minimal model without actin turnover

To keep the discussion straightforward, we first neglect actin turnover dynamics, which amounts to assuming fast flows ($L/v_0 <$ turnover time, of the order of minutes). The actin cortex can then be simply modelled as a viscous incompressible fluid. Force balance in the fluid and incompressibility then make it possible to compute exactly the actin flow profile $v_x, v_y$ in the cortex, as well as the pressure field $p$ in the cortex in the limit of small deformation ($\epsilon \ll 1$); see last section for an explicit derivation. The $x$ component of the resulting force applied by the wavy walls on the cell is then deduced by balancing the normal stress in the cortex, and yields after integration over the surface of the two wavy confining walls in contact with the cell,

$$F_x = -\frac{8\pi^6}{3}\frac{z_0 L v_0 \eta}{\delta}\left(\frac{\delta}{\lambda}\right)^6 \epsilon^2, \tag{1}$$

where we recall $\epsilon = 2w_1/\delta$. The force is therefore opposed to the direction of the actin flow (recall that $v_0 < 0$) and propels the cell forward. The dependence on the actin viscosity shows that motion is induced by shear forces, and crucially controlled by the roughness of the geometry (parametrised by the wave length $\lambda$ and amplitude $w_1$ of the wall deformation). Note that this force is induced by an asymmetry of the stress in the actin cortex layer ; an additional hydrostatic pressure in the cytoplasm could be taken into account, but does not contribute to motion after integration over the cell surface. It is at this stage instructive to give orders of magnitude. For classical values $\eta \sim 10^5$ Pa.s see[18] and $v_0 \sim 1 - 10$ $\mu$m/min see[19], one finds for realistic geometries used in experiments, for which $\lambda$ ranges from 5 to 25 $\mu$ m, the channel depth $z_0 \sim 10\mu$m, cell length $L \sim 20$ $\mu$m, and $\epsilon = w_1/h_0 \sim 1$, we find forces in the range $0.1 - 100$ nN, which is an expected order of magnitude for migrating cells[17].

In the general case, where $L$ isn't a multiple of $\lambda$, we can still express the total force as an integral over the contact region between the flowing layer and the channel from $x = 0$ to $x = L$. Then, the total force is the integer part of $L/\lambda$, call it $n$, times $F_x$, plus some residual integral over less than a period that may be positive or negative depending of relative position of the cell with the wavy pattern of the channel. This residual is negligible when $L \gg \lambda$, but for finite $L/\lambda$ could be of order $F_x$, and depends on the relative position of the cell in the channel, thereby resulting in start-stop-type motion, and eventually cell arrest when $L$ is of the order of $\lambda$. For example, if $L = \lambda/2$, it is possible that the "cell" is only in contact with backward-facing channel walls, which means that migration is impossible.

##### B. Effect of actin turnover

Actin turnover can be taken into account in the above model in order to give a more realistic description of the actin cortex. A classical model consists in assuming that F-actin is polymerised at the membrane, yielding an inward local speed $v_p$ of the actin normal to the cell/wall interface. In oder to reach a steady state a depolymerisation rate $k_d$ must be introduced. In the case of a flat interface, this simple model yields a stationary cortex of constant thickness $\delta_0 = v_p/k_d$. In the presence of a wavy wall as depicted in Fig. 3, and assuming an average cortical flow along the $x$ axis of speed $v_0$, the full actin flow profile and pressure fields can be obtained using similar methods. This analysis yields the same propulsion force (up to a numerical prefactor)

$$F_x \sim \frac{z_0 L v_0 \eta}{\delta_0} \left(\frac{\delta_0}{\lambda}\right)^6 \epsilon^2, \tag{2}$$

as in the case without turnover, with however a cortex thickness $\delta_0$ imposed by the actin dynamics. This shows the robustness of the results.

##### C. Connection between propulsion force and cell velocity

Our model makes predictions on the dependence of the cell propulsion force on the channel shape. The connection between channel shape and cell velocity, however, is less clear, and we caution that the predicted scaling of the propulsion force with channel wavelength does not necessarily apply directly to the cell velocity. On general grounds, one can argue that the cell speed $U$ is related to $F_x$ by the relation $\xi U = F_x$, where $\xi$ is an effective drag coefficient due, for example, to the work needed to move and deform the nucleus [20]. There are however a number reasons to expect a non-trivial relationship between cell velocity and channel shape parameters ($\lambda$ and $w_1$). First, we note that the model assumes that $v_0$ is the actin flow speed with respect to the *substrate*, and is prescribed. In all likelihood, $v_0$ is regulated by the cell and can depend on the geometry of the environment — in a similar way that it depends on adhesion conditions[21]. Second, the effective friction $\xi$ also depends on the channel shape (that is, $\lambda$ and $w_1$); its functional dependence is not clear and would need further assumptions to be modelled. Third, since, experimentally, we are not in the regime $L \gg \lambda$, the residual force introduced above comes into play, so that the exact proportionality to the explicit expression $F_x$ is lost. Last, the model describes the actin gel as a continuous viscous fluid; such description is valid at scales sufficiently larger that the gel mesh size. A different behaviour is therefore expected for geometric irregularities at small length scales ($\lambda < 0.1 - 1\mu$m). For all these reasons, we cannot conclude that the resulting cell speed scales quantitatively the same way as the predicted propulsion force for fixed $L$ and $v_0$.

#### III. DERIVATION OF THE RESULTS

In this section, we derive the main results given in Eq. 1. As justified above, we suppose that the actin cortex can be described as an incompressible viscous fluid. Furthermore, at the cell scale we can neglect fluid inertia. The fluid thus satisfies the Stokes equations and the incompressibility constraint:

$$\partial_x p = \eta \left(\partial_x^2 + \partial_y^2\right) v_x \tag{3a}$$

$$\partial_y p = \eta \left(\partial_x^2 + \partial_y^2\right) v_y \tag{3b}$$

and

$$\partial_x v_x + \partial_y v_y = 0 \,, \tag{4}$$

where $\partial_x \equiv \frac{\partial}{\partial x}$ and $\partial_y \equiv \frac{\partial}{\partial y}$. Here $p$ is the pressure, $\eta$ is the fluid viscosity, and $v_x$ and $v_y$ are the components of the velocity field $\mathbf{v}$. The problem is thus rephrased as a rather classical question of stokes hydrodynamics in undulated geometry (see for example[22]), which we adapt here to the appropriate geometry of cortical dynamics.

For flat channel walls ($\epsilon = 0$) with perfect slip (zero adhesion), the fluid undergoes "plug flow" with velocity components $v_x = v_0$ and $v_y = 0$. We then perturb around this $\epsilon = 0$ solution. Thus, to $O(\epsilon)$ we write

$$v_x(x, y) = v_0 + \epsilon \sin(kx) f'(y) \tag{5}$$

$$v_y(x, y) = -k\epsilon \cos(kx) f(y) \,, \tag{6}$$

where the function $f(y)$ is to be determined. Note that incompressibility is automatically satisfied by this Ansatz. By symmetry of both the channel geometry and boundary conditions, it follows that $f(y)$ is an even function. Similarly, we write down an Ansatz for the pressure, valid to $O(\epsilon)$:

$$p(x, y) = \epsilon \cos(kx) g(y) \,. \tag{7}$$

Since $\nabla^2 p = 0$, we immediately obtain $g(y)$, which is an odd function, as

$$g(y) = A \sinh(ky) \,, \tag{8}$$

with $A$ an unknown constant. Thus, from Eq. (3), $f$ satisfies

$$f''(y) - k^2 f(y) = -\eta^{-1} A \cosh(ky). \tag{9}$$

Two boundary conditions are then needed to determine the even function $f(y)$. First, no-flux across both the channel wall and the cortex-cytoplasm interface means $\mathbf{v}\cdot\hat{\mathbf{n}}|_{y=\pm h_0} = (v_y - h'_\pm\, v_x)\big|_{y=\pm h_0} = 0$. Noting that $h'_- = h'_+ = h_0\epsilon k\cos(kx)$, this implies

$$f(y = h_0) = -v_0\,. \tag{10}$$

Second, the no-friction condition at the wall and cortex-cytoplasm interface is equivalent to vanishing of the shear stress at these locations. Denoting the total stress tensor in the fluid as $\sigma - p\mathbf{1}$, with $\mathbf{1}$ the unit tensor, the viscous part of the stress is given by $\sigma$. The shear stress at an interface is given by $\sigma_{tn} = \hat{\mathbf{t}}\cdot(\sigma|_{y=\pm h_0})\cdot\hat{\mathbf{n}}$, where $\hat{\mathbf{t}} \simeq (1, h'_\pm)$ and $\hat{\mathbf{n}} \simeq (\mp h'_\pm, 1)$ are the local unit vectors tangent and normal to the interface, with the normal pointing away from the fluid. To leading order in $\epsilon$, $\sigma_{tn} \simeq \sigma_{xy}|_{y=\pm h_0}$, and thus the second boundary condition boils down to $\sigma_{xy}|_{y=\pm h_0} = 0$. This condition leads to

$$f''(y = h_0) = k^2 v_0\,, \tag{11}$$

where we have used $\sigma_{xy} = \eta\,(\partial_x v_y + \partial_y v_x) = \eta\epsilon\,(f'' + k^2 f)$ and $f(y = h_0) = -v_0$. Thus,

$$f(y) = -\frac{A}{2k\eta}y\sinh(ky) + \frac{B}{\eta k^2}\cosh(ky)\,, \tag{12}$$

where

$$A = -2\eta h_0 k^2 v_0/\cosh(h_0 k) \tag{13}$$

$$B = -\eta h_0 k^2 v_0\ \left(h_0 k\tanh(h_0 k) + 1\right)/\cosh(h_0 k)\ , \tag{14}$$

similar to the result found in[22]]. Finally, the force per unit area exerted by the walls along $x$ is simply the total stress tensor at $y = h_+$, projected along the unit vector $\hat{\mathbf{n}}$ and along $\hat{\mathbf{x}}$. The total force is then obtained after integration over the surface of the wavy walls, $S_0$, in contact with the cell. Assuming that the cell is confined between two identical wavy walls, this gives:

$$\begin{aligned} F_x &= \int_{S_0} dS\ \hat{\mathbf{x}}\cdot(\sigma - p\mathbf{1})|_{y=h}\cdot\hat{\mathbf{n}} \\ &\simeq -2z_0\int_0^L dx\ h'_+(x)\ \left.(\sigma_{yy} - p)\right|_{y=h_0} \\ &= -4z_0 L\eta k^3 h_0^2 v_0\epsilon^2\frac{[\sinh(h_0 k)\cosh(h_0 k) - h_0 k]}{\cosh(2h_0 k) + 1}\,, \end{aligned} \tag{15}$$

where we have used $\sigma_{yy} = 2\eta\partial_y v_y$, and having calculated $p$ and $v_y$ using Eqs. (7), (8), and (12)-(14). Here we have assumed that $L$ is an integer multiple of the wave length $\lambda$. Making use of the cortex width $\delta$ and wave length $\lambda$, this takes the simpler form in the limit $h_0 k \to 0$ (long wavelengths):

$$F_x = -\frac{8\pi^6}{3}\frac{z_0 L v_0 \eta}{\delta}\left(\frac{\delta}{\lambda}\right)^6 \epsilon^2, \tag{16}$$

which gives the main dependence on parameters discussed above, where we recall $\epsilon = 2w_1/\delta$.

Finally, we note that the propulsion force, $F_x$, is second order in the channel modulation amplitude, $w_1$ (or equivalently $\epsilon$). This means that, in order to satisfy force balance along $x$ for a cortical element, a subleading $O(\epsilon^2)$ correction should be included in the cortical pressure $p$. This higher order contribution, averaged over a period, leads to a constant pressure gradient in the cortex that can be written

$$\nabla_x p = -\frac{4\pi^6}{3}\frac{v_0 \eta}{\delta^2}\left(\frac{\delta}{\lambda}\right)^6 \epsilon^2. \tag{17}$$

This contribution to the stress, however, does not affect our leading order determination of $F_x$.
