## Supplementary Data for "Adhesion-free cell migration by topography-based force transduction"

**This PDF file includes:**

Materials and Methods
Supplementary Table S1
Extended Data Fig. 1 and 2
Captions for Extended Data Fig. 1 and 2
Captions for supplementary videos 1 to 10

**Other Supplementary Materials for this manuscript includes the following:**

Supplementary Text: Model
Supplementary videos 1 to 10

Materials and Methods

**Cell culture.** Cells were grown and maintained in a humidified incubator at 37ºC and 5% CO2. LMR7.5 T cell hybridoma, gift from A.M. Lennon-Duménil (Institut Curie, Paris, France), were cultivated in R10 medium (RPMI 1640 supplemented with 10% fetal bovine serum, 2 mM L-glutamine, 100 units/ml penicillin, 100 µg/ml streptomycin, and 50 µM 2-mercaptoethanol; Gibco, Thermo Fisher Scientific, USA). Lifeact-eGFP [23] parental T cell line was generated by nucleofection with eGFP reporter construct (Kit V, Lonza, Switzerland). Alternatively, LMR T cells were infected with a plasmid coding for Lifeact-mCherry (pLenti6.3, Invitrogen, Thermo Fisher Scientific, USA). *LX-293 HEK cells* (Clontech, USA) were used for lentivirus production, and maintained in D10 (DMEM with 2 mM L-glutamine, 10% fetal bovine serum; 100 units/ml penicillin and 100 µg/ml streptomycin; Gibco). All cell lines were frequently tested and free of mycoplasma.

**Reporter and CRISPR/Cas9 cell line generation.** ***CRISPR/Cas9 design and production***. Single guide RNA designs were performed as described elsewhere [24]. Briefly, single guide RNAs were designed to induce the double strand break in the first exons of the mice talin1 gene. Then sgRNAs were scored for high on-target effects (https://portals.broadinstitute.org/ gpp/public/analysis-tools/sgrna-design), and low off-target (http://crispr.mit.edu). sgRNAs were cloned into LentiCRISPRv1 vector (a gift from Feng Zhang, Addgene plasmid #49535 and sequenced. Guides sequences are: sgRNA-Scramble s(5' -3') GCCGTGGCGCATGGGTAGCA; sgRNA-Talin1g1 s(5' -3') ATAATGCCCTACGAGCCGT (off-target score 91); sgRNA-Talin1g 2 s(5' -3') CTCACTGTTTCCCCGGGTA (off-target score 82); sgRNA-Talin1g 3 s(5' -3') GTCGAGGCTGGGCGACTGG (off-target score 98). ***Lentivirus production*** was performed as described in [25]. Briefly, LX-293 HEK cells (Clontech, USA) were co-transfected with LentiCRISPRv1- or LentiCRISPRv2-, packaging- psPAX2 (AddGene 12260) and pCMV-VSV-G envelope- plasmids using Lipofectamine 2000 (Thermo Fisher Scientific, USA) as recommended by the manufacturer, and cells were resuspended in R10 one day prior transfection. The surpernatant was collected after 72 h and stored at -80℃. ***CRISPR-based talin KO generation.*** Lifeact-eGFP LMR7.5 T cells were spin-infected at 1500g for 1 h in the presence of the lentivirus-containing supernatant and 6µg/ml Polybrene

(Sigma-Aldrich, USA). Spin infection was carried out in a 12-well plate with 3x10^5 cells/ml per reaction plus 500µl of undiluted virus. Three days after infection, cells were selected for stable virus insertion using 5µg/ml Puromycin (Gibco, Thermo Fisher Scientific, USA) for 7 days and were subjected to western blotting. Cells were then sorted and expanded into clones in puromycin-containing R10 using fluorescence-activated cell sorting (FACS Aria III, BD Biosciences). *T cell infection and selection.* For infection of LMR7.5 with Lifeact-mCherry reporter construct, the same infection protocol was applied except cells were selected with 5µg/ml Blasticidin (Gibco, Thermo Fisher Scientific, USA) and sorted for fluorescence expression.

**Western blot.** For immunodetection of Talin and GAPDH proteins, 1x10^6 control and talin KO cells were centrifuged and lysed in RIPA buffer (New England Biolabs, USA) and supernatant was denaturated in 2X Laemmli Buffer (161-0737, Biorad, ) at 95°C for 5 minutes. Protein lysates where loaded on a 10% tris-glycine gel (Invitrogen, Thermo Fisher Scientific, USA), separated by SDS-PAGE and further transferred by electrophoresis to a polyvinylidene difluoride membrane (iBlot, Invitrogen, Thermo Fisher Scientific, USA) according to the manufacturers instructions, with a 9 min transfer time. Membranes were blocked in PBST-BSA 5% for 1 h and incubated with the primary antibodies overnight at 4°C. Antibodies against Talin (mouse anti-pan-talin, clone 8d4, T3287, Sigma-Aldrich, USA) and GAPDH (mouse, clone GA1R, ab125247, Abcam, UK) were diluted 1/200 and 1/3000 in PBST-BSA 1%, respectively. After washing with TBST, the secondary antibody was applied for 1 hour in PBST-BSA 5% at room temperature (goat anti-mouse IgG HRP conjugate, diluted 1:5000, 170-6516, Biorad, USA). Protein detection was performed by enhanced chemoluminescence (ECL, Thermo Fisher Scientific, USA) detection using a VersaDoc imaging system (Biorad, USA).

**Adhesion assay**. Control and talin KO T cell (5x10^5) were plated in pre-warmed R10 medium in a 25 $cm^2$ culture flask (TPP, Sigma-Aldrich, USA), and placed in a humidified incubator at 37°C, 5% $CO_2$, 24 h before observation. Cells were imaged with an inverted microscope at low magnification (DM IL Led and DFC450 digital camera, Leica Microsystems, Germany) and quantification of the cells adhering to the bottom of the dish was performed. The experiment was carried out 4 times.

**Spreading assay.** 2D spreading assay was performed on glass coverslip (Menzel-Glaser #1, VWR, USA) that were previously coated on a small spot with a mix of 1µg of anti-CD3e antibody (Clone 145-0281-85, 16-0031-85, eBioscience, Thermo Fisher Scientific, USA) and 1µg of anti-CD28 antibody (clone 37.51, 16-0281-85, eBioscience, Thermo Fisher Scientific, USA) overnight at 4ºC, and washed 3 times with PBS. 0.2x10^6 control (LifeAct-mCherry) and talin KO (LifeAct-GFP) T Cells were gently mixed in 200µl R10 and allowed to settle on the coated dish for 60 min at 37ºC and 4.5%CO2. Cell spreading was then imaged by simultaneous TIRF- and epifluorescence microscopy.

**Photolithography.** Photolithography was performed as described in [26] and [27] with the following adaptations. ***Photomasks.*** Patterns were designed with Coreldraw X8 (Corel Corporation, USA) exported to DXF, then converted to GERBER format with LinkCad. 100 mm or 125 mm chrome photomasks (PhotoData/JD Photo-Tools, UK) were used. ***Confiners.*** 3 µm confiner masters were made by spin coating SU8-GM1050 (Gersteltec, Switzerland) for 40s at 3000 RPM, prebaking for 1 min at 120°C, exposing 35 mJ/cm² of UV, post exposure baking for 5 min at 95°C, developing in SU8 developer, and then hard baking at 150°C for 5 min. The 5 µm confiner master was made by spin coating SU8-2005 (Microchem, USA) for 30s at 3000 RPM, prebaking 2 min at 95°C, exposing 105 mJ/cm² of UV, post exposure baking for 3 min at 95°C, developing in SU8 developer, and then hard baking at 150°C for 5 min. ***EDTA device.*** A 4 µm high master was made by spin coating SU8-2005 for 30 seconds at 5000 RPM, prebaking 2 min at 95°C, UV exposure of 550 mJ/cm² through a PL- 360-LP (Omega Optical) optical filter on an EVG mask aligner 610 (EVG group, Austria). The wafer was post exposure baked for 3 min at 95°C, developed in SU8 developer, and then hard baked at 150°C for 5 min. The optical filter was necessary for high resolution features in SU8. ***Multilayer devices.*** (1) The 25 µm high control master was made by spin coating SU8-3025 (Microchem, USA) for 30 seconds at 3000 RPM. The wafer was soft baked for 15 min at 95°C, then exposed to 200 mJ/cm² UV, then post exposure baked at 95°C for 5 min, then developed in SU8 developer. (2) The fluid layer master was first fully coated with a submicron layer of GM1040-SU8 by spin coating at 5000 RPM for 2 min. The wafer was then baked at 95°C for 5 min, exposed to 100 mJ/cm² UV, post exposure baked at 95°C for 30 min, and then developed in SU8 developer. The 5 µm fluid layer features were made on top of the coated wafer by spin coating SU-2005 for 30 seconds at 3000 RPM, soft baking for 2 min, UV exposure of 550 mJ/cm² through the optical filter, post exposure baking for 3 min at 95°C, and developing in SU8 Developer. (3) The fluid layer master was then prepared for the next layer by spin coating HMDS at 3000 RPM for 30 seconds, and baking 1 min at 126°C. Scotch tape was applied over the alignment marks. AZ-40XT-11D (MicroResist Technologies, Germany) was then coated at 3000 RPM for 30 seconds. The scotch tape was then peeled off to reveal the alignment marks. The wafer was then baked at 126°C 7 min, aligned, exposed to UV 500 mJ/cm². After post-exposure bake 10 min at 120°C, it was developed in 726 MIF for about 5-10 min. The features were then rounded to make parabolic channels at 130C for 2 min (26.6 µm – center height). ***Silanization.*** Before applying PDMS, wafers were placed in a desiccator with 10 µL of Trichloro(1H,1H,2H,2H-perfluorooctyl)-silane (Sigma-Aldrich, USA), a vacuum of 100 mbar was applied, and then the desiccator was sealed for 1 hour. One permanent coating was sufficient for later uses.

**PDMS devices microfabrication.** Microfabricated devices were generated as previously described [11,26,27], with 1:10 PDMS and degased 2 min at 2000 RPM (mix) and 2 min at 2200 RPM (defoam) in a Mixer/defoamer (ARE-250, Thinky, USA) prior to use. ***Confiner.*** PDMS (Sylgard 184, Ellsworth Adhesives, USA) was gently poured onto the wafers, then 10 mm coverslips were activated by plasma cleaning 2 min at medium intensity (Harrick Plasma Cleaner, pdc-002, Harrick Plasma, USA) and pressed upside-down onto the PDMS-covered wafers. The wafer was baked on a hot plate at 95ºC for 15 min, and the 300 µm-width micropillars-coated coverslip were gently removed from the wafer. A soft PDMS 10 mm-diameter pillar (1:30) was poured into a home-made metal mould, degassed under vaccum and baked at 80ºC for a week. ***EDTA device.*** 20-25g of 1:10 PDMS (Sylgard 184, Ellsworth Adhesives, USA) was poured onto the wafer contained in an aluminium mould in a petri dish, degased in a vacuum desiccator and baked overnight at 80ºC. The devices were then diced with a razor blade and 2 mm entry/exit holes were punched (Harris Unicore biopsy puncher, Sigma-Aldrich, USA). Following punching, they were cleaned with tape, sonicated in ethanol, blown dry and plasma bonded to a coverslip. ***Multilayer devices.*** 80g of PDMS (RTV615, Techsil, UK) was used; 10g of PDMS was put on the control layer and spin coated (500 RPM for 15 s followed by 2300 RPM for 60 s), and 70g was applied on top of the flow layer in a tight aluminium mould and further degased in a vacuum desiccator to pump out the remaining bubbles. Both layers were baked at 80ºC for 45 min, and the flow layer was removed from the mould, trimmed with a razor blade, and entry and exit holes were punched with a home-made arbor press. Debris from both the flow device and the control layer on its wafer were carefully removed with scotch tape, the 2 layers were plasma cleaned (2 min at medium intensity), aligned manually and bonded together. After overnight baking in 80ºC, the chip was removed from the wafer, the control layer entry holes were punched, and the chip was bonded to the glass coverslip by oxygen plasma, and kept at 80ºC until further use. Before the experiments, all entry ports were connected with metal pins (NE-13-1003, New England Small Tube Corporation, USA).

**Migration assays.** Before experiments, 2x10^5 T cells in 2 ml R10 were stained with Hoechst 33342 for 30 min (1 drop, NucBlue, R37605, Invitrogen, Thermo Fisher Scientific, USA) when nucleus visualization was needed for cell tracking. ***Collagen assay.*** 3D collagen scaffold was obtained by mixing bovine collagen (PureCol, Advanced BioMatrix, USA) in 1X minimum essential medium eagle and 0,4% sodium bicarbonate (both Sigma-Aldrich, USA), with 3x10^5 cells in R10 at a 2:1 ratio [5,28]. After casting the mix in home-made migration chambers, gels were allowed to polymerize 45 min at 37ºC 5%$CO_2$. Migration was observed by time-laps videomicroscopy and analysed with Trackmate. ***Cell confiner assay.*** PDMS micropillars were placed on the soft pillar on a home-made magnetic glass lid. The set-up was placed in a dish containing R10 in a humidified incubator at 37ºC 5% $CO_2$ 1 h prior to experiments. Cells (5x10^4 in 5 µl R10) were then pipetted onto the medium-freed micropillars and confined on a dish above a magnetic ring, and R10 was added back in the dish ready for imaging. ***EDTA migration assay.*** T cells (5x10^4) were introduced into the 2 mm entry port of the microchamber, and the microchamber was soaked in R10. Cells were left to enter freely into the confinement and channel zone for 1 hour, and then imaged with a videomicroscope. After 1 h of imaging, chemical disruption of the cell adhesions was performed by removing the R10 from the dish and adding R10 containing 10 mM EDTA (prepared from 0,5M stock, EDS-100G, Sigma-Aldrich, USA). Imaging was then resumed for 1 h. ***Multilayered***

***microfluidics: maze and microchannels migration assay.*** As talin KO failed to freely enter the confinement zone, a home-made microfluidic set-up was used to push the cells by a applying a pressure differential to the in- and outlet ports of the device. Microfluidics designs in Extended Data Fig. 2a were used, as well as separated chambers containing the 4 pillar mazes (identical to [26] with 4 chambers). Briefly, the core component of those devices are the flow layer with 5 µm height pillar maze (Fig. 2a) or channels (Fig. 2h, 2j and Fig. 3a) with an exit, cell and medium entry ports and on one side and on the other side a sink channel connected to 2 exits. Flow and sink channels are rounded, 26 µm high. For controlled flow of cells and fluids, all ports are equipped with independently controllable push-up PDMS membrane valves. Briefly, the control channels are connected to solenoid valves (MH1, miniature, Festo, USA) controlled with a Matlab graphical user interface (MathWorks, USA). First the chip was mounted into the 37ºC 5% $CO_2$ chamber of the microscope, and optimal closing pressures of 0,15-0,2MPa of the water-filler PDMS membrane valves were determined for each chip, each valve checked individually by microscopy. Second, the whole chip was saturated with pre-warmed medium by applying a pressure of 0.04MPa, and incubated in R10 for 30 min. Cells were first loaded via the ports on one side, resulting in their distribution along the chamber, then slowly pushed with a pressure <0.01Mpa from the medium port toward the confinement zone, and further pushed in the microchannels or micropillars maze. Finally, all valves were closed to ensure that no external pressure or flow was applied to the cells during imaging. ***TIRF microfluidics: microchannels migration assay.*** For Fig. 1e-g, TIRF microscopy was not possible with the multi-layered microfluidic devices, so devices with only the flow layer were generated. First R10 medium was forced into the device with a vacuum desiccator, and the chips were incubated in humidified 5% $CO_2$, 37 °C, for 2 h prior to experiments. Experiments were performed on the microscope stage (humidified environment 5% $CO_2$, 37 °C) and a low cell flow was delivered and stopped after cell loading in the microchannel using a LA120 syringe pump (5 µl/hour, Landgraf Laborsysteme, Germany).

**Time-lapse videomicroscopy.** Brightfield movies of T cells for collagen assays (10X objective) and 3 µm-high confiner assays (20X objective) were performed by time-lapse acquisition (time interval 20s) using inverted cell culture microscopes (DM IL Led, Leica Microsystems, Germany) equipped with cameras (ECO415MVGE, SVS-Vistek, Germany) and custom-built climate chambers (5% $CO_2$, 37 °C, humidified). 5 µm-high confiner, EDTA and microfluidics assays were recorded every 30 or 60 seconds and 2 to 6 multi-positions with NIS Elements software (Nikon Instruments, Japan). Experiments were performed with an inverted widefield Nikon Eclipse Ti microscope in a humidified and heated chamber at 37°C and 5% $CO_2$ (Ibidi Gas Mixer), equipped with a 20x/0.5NA PH1 air objective, a Hamamatsu EMCCD C9100 camera and a lumencor light source (390nm, 475nm, 542/575nm; Nikon Instruments, Japan). Total internal reflection (TIRF) microscopy was performed with a 60X/1.46NA oil objective, optovar 1X or 1.6X in a humidified and heated chamber at 37°C and 5% $CO_2$ using an inverted Axiovert 200 (Zeiss, Germany) microscope, a TIRF 488/561-nm laser (Visitron systems, Germany) and an Evolve EMCCD camera (Photometrics, USA) triggered by VisiView software (Visitron systems, Germany). Assays were recorded every second for TIRF acquisition only (Fig. 1e-g, supplementary video 3 example 3) or every 2 or 3 seconds when acquisition of both TIRF and brighfield was necessary (Fig. 4e-g, Extended Data Fig. 2e-g, supplementary video 3 example 1 and 2, and supplementary video 10). When cells migrate outside the field of view (supplementary video 10), they were manually followed via a joystick controller.

**Image Analysis.** FIJI imaging processing software (https://fiji.sc) was used for image and videomicroscopy analysis. ***Spreading assay.*** TIRF images of single cells were binarized after de-speckling and background subtraction, and spreading area was measured using the Analyse Particles tool. ***Manual Tracking.*** 1 h-long brightfield movies of T cells in collagen and 3 µm-high confiner were reduced to a 1 and 2 min interval, respectively, and cells were tracked manually by using the 'Manuel tracking' plugin provided by Fiji. ***Automated Tracking.*** For the 5 µm-high confiner, pillar maze and EDTA assays, cell migration was analysed by nucleus tracking using Trackmate [29] (https://imagej.net/TrackMate), resulting in higher cell speed due to nucleus shifting inside the cell (Fig. 1d, supplementary video 2). To reduce the wobbling effect, directionality was calculated from Trackmate files with a home-made script (Matlab, MathWorks, USA) by dividing the displacement of the cell

(Euclidian distance) by its total path length (Extended Data Fig. 1g). For the EDTA assay, tracks before and after EDTA treatment were analysed and the speed was extracted at the single cell-level in the different zones (2,5D confinement, smooth and serrated microchannels containing only one cell). For Fig. 4b, cell migration in the channel zones was regarded as efficient for velocities above 0.5 µm/min. ***Actin flow vs cell velocity analysis***. For Fig. 1e-g, Fig. 4e-g and Extended Data Fig. 2e-g, kymographs were generated along the cells trajectories, and both cell and actin velocities were extracted from those kymographs. To follow cells migrating in a channel with a high magnification (Fig. 4e-g and supplementary video 10), the channel were re-aligned with a home-made script (Matlab, MathWorks, USA). ***Microchannels assays analysis***. Cells going in both directions of the microchannels were analysed, and channels containing more than one cell were excluded. Kymographs were generated along the channel, and cell velocity was calculated for each cell in the different zones.

**Statistical analysis.** All statistical analyses were performed with GraphPad Prism (GraphPad Software, USA). See Supplementary Table for exhaustive experiments and analytical summary. For graphs in Fig. 1, Fig. 2g, 2i and 2l, non parametric T test (Mann-Whitney, 2 tailed, 95% confidence interval) was performed. For Fig. 2l, non parametric paired T test was performed (Wilcoxon matched-pairs signed rank test). For Fig. 2d, 2e, 2k, Fig. 3d and Extended Data Fig. 2c, one-way ANOVA with Kruskal-Wallis test (with Dunn's post-test) was used. For Fig. 3g one-way ANOVA with Friedman test (Dunn's post-test) was performed. For box and whiskers graphs in Fig. 4b, 4g and Extended Data Fig. 2c the box extends from the 25th to the 75th percentiles, with the middle line showing the median, and the whiskers show min to max values.

**Data availability.** All data are available from the authors on request.

**Materials and Methods References**

11. Le Berre, M., Aubertin, J. & Piel, M. Fine control of nuclear confinement identifies a threshold deformation leading to lamina rupture and induction of

specific genes. *Integr Biol* **4,** 1406–1414 (2012).
23. Riedl, J. *et al.* Lifeact: a versatile marker to visualize F-actin. *Nat Methods* **5,** 605–607 (2008).
24. Shalem, O. *et al.* Genome-Scale CRISPR-Cas9 Knockout Screening in Human Cells. *Science (80-. ).* **343,** 84–87 (2014).
25. Leithner, A. *et al.* Diversified actin protrusions promote environmental exploration but are dispensable for locomotion of leukocytes. *Nat. Cell Biol.* **18,** 1253–1259 (2016).
26. Leithner, A., Merrin, J., Reversat, A. & Sixt, M. Geometrically complex microfluidic devices for the study of cell migration. (2016). at <http://dx.doi.org/10.1038/protex.2016.063>
27. Schwarz, J. *et al.* A microfluidic device for measuring cell migration towards substrate-bound and soluble chemokine gradients. *Sci. Rep.* (2016). doi:10.1038/srep36440
28. Sixt, M. & Lämmermann, T. in *Cell Migration: Developmental Methods and Protocols* (eds. Wells, C. M. & Parsons, M.) 149–165 (Humana Press, 2011). doi:10.1007/978-1-61779-207-6_11
29. Tinevez, J. Y. *et al.* TrackMate: An open and extensible platform for single-particle tracking. *Methods* **115,** 80–90 (2017).

Supplementary Table S1

| Assay | Figure | Number of experiments | Number of cells: Control | Number of cells: Talin KO | Columns | Statistical test | Significance | P-value |
|---|---|---|---|---|---|---|---|---|
| 2D Culture | 1a | 4 | na | na | na | T test. Paired, 2 tailed. 95% confidence interval | *** | 0.00076 |
| 2D Spreading | 1b | 2 | 139 | 81 | na | T test. non parametric. Mann-Whitney. 2 tailed. 95% confidence interval | **** | <0.0001 |
| 3D Collagen | 1c | 3 | 51 | 52 | na | T test. non parametric. Mann-Whitney. 2 tailed. 95% confidence interval | n.s. | 0.628 |
| 2,5D 5µm Confiner | 1d | 4 | 290 | 284 | na | T test. non parametric. Mann-Whitney. 2 tailed. 95% confidence interval | **** | <0.0001 |
| 2,5D 5µm Confiner TIRF | 1g | 3 | 43 | 37 | na | T test. non parametric. Mann-Whitney. 2 tailed. 95% confidence interval | **** | <0.0001 |
| | | | | | na | T test. non parametric. Mann-Whitney. 2 tailed. 95% confidence interval | **** | <0.0001 |
| Maze Control – Speed | 2d | 3 | 88 | na | 3µm vs. 4µm | One-way ANOVA with Kruskal-Wallis test with Dunn's post-test | ns | >0.9999 |
| | | | | | 3µm vs. 5µm | | ns | 0.4238 |
| | | | | | 3µm vs. 6µm | | ns | >0.9999 |
| | | | | | 4µm vs. 5µm | | ns | 0.2019 |
| | | | | | 4µm vs. 6µm | | ns | >0.9999 |
| | | | | | 5µm vs. 6µm | | ns | 0.5959 |
| Maze Control – Displacement | 2E | | | | 3µm vs. 4µm | One-way ANOVA with Kruskal-Wallis test with Dunn's post-test | ns | >0.9999 |
| | | | | | 3µm vs. 5µm | | ns | 0.1162 |
| | | | | | 3µm vs. 6µm | | ns | 0.6308 |
| | | | | | 4µm vs. 5µm | | ns | >0.9999 |
| | | | | | 4µm vs. 6µm | | ns | >0.9999 |
| | | | | | 5µm vs. 6µm | | ns | >0.9999 |
| Maze TalinKO – Speed | 2d | 6 | na | 79 | 3µm vs. 4µm | One-way ANOVA with Kruskal-Wallis test with Dunn's post-test | ns | 0.1282 |
| | | | | | 3µm vs. 5µm | | ns | >0.9999 |
| | | | | | 3µm vs. 6µm | | ns | >0.9999 |
| | | | | | 4µm vs. 5µm | | ns | 0.1220 |
| | | | | | 4µm vs. 6µm | | ** | 0.0021 |
| | | | | | 5µm vs. 6µm | | ns | 0.8944 |
| Maze TalinKO – Displacement | 2E | | | | 3µm vs. 4µm | One-way ANOVA with Kruskal-Wallis test with Dunn's post-test | ns | 0.8168 |
| | | | | | 3µm vs. 5µm | | ns | >0.9999 |
| | | | | | 3µm vs. 6µm | | ns | 0.0814 |
| | | | | | 4µm vs. 5µm | | ** | 0.0089 |
| | | | | | 4µm vs. 6µm | | **** | <0.0001 |
| | | | | | 5µm vs. 6µm | | ns | 0.4732 |
| 2,5D 3µm Confiner | 2g | 4 | 52 | 60 | na | T test. non parametric. Mann-Whitney. 2 tailed. 95% confidence interval | **** | <0.0001 |
| Channel Smooth | 2i | | 99 | 24 | na | T test. non parametric. Mann-Whitney. 2 tailed. 95% confidence interval | **** | <0.0001 |
| Channel Serrated | 2k | 3 for Control, 4 for Talin KO | Serrated zone / Smooth zone | Serrated zone / Smooth zone | Control - Serrated zone vs. Control - Smooth zone | one-way ANOVA with Kruskal-Wallis test with Dunn's post-test | ns | >0.9999 |
| | | | 57 / 39 | 8 / 5 | Control - Serrated zone vs. Talin KO - Serrated zone | | ns | 0.3313 |
| | | | | | Control - Serrated zone vs. Talin KO - Smooth zone | | *** | 0.0009 |
| | | | | | Control - Smooth zone vs. Talin KO - Serrated zone | | ns | 0.6685 |
| | | | | | Control - Smooth zone vs. Talin KO - Smooth zone | | ** | 0.0028 |
| | | | | | Talin KO - Serrated zone vs. Talin KO - Smooth zone | | ns | 0.4040 |
| EDTA | 2l-m | 3 | 73 | na | Confinement vs Confinement+EDTA | Wilcoxon matched-pairs signed rank test, two tailed | **** | <0.0001 |
| | | | | | | Pairing : rs (Spearman) 0,4372 | **** | <0.0001 |
| | | | 23 | na | Serrated Channel vs Serrated Channel+EDTA | Wilcoxon matched-pairs signed rank test, two tailed | * | 0.0167 |
| | | | | | | Pairing : rs (Spearman) 0,6212 | *** | 0.0008 |
| | | | 27 | na | Smooth Channel vs Smooth Channel+EDTA | Wilcoxon matched-pairs signed rank test, two tailed | **** | <0.0001 |
| | | | | | | Pairing : rs (Spearman) 0,3756 | * | 0.0268 |
| Channel variable complexity | 4b | 3 | na | Migration / No Migration | | Fisher's exact test, two sided | **** | <0.0001 |
| | | | | 37 / 4 | 6-12µm | | | |
| | | | | 5 / 17 | 24µm-Smooth | | | |
| Channel variable complexity – Control | 4d | 4 | 88 | na | 6µm vs. 12µm | One-way ANOVA with Kruskal-Wallis test with Dunn's post-test | ns | >0.9999 |
| | | | | | 6µm vs. 24µm | | ns | >0.9999 |
| | | | | | 6µm vs. Smooth | | * | 0.0234 |
| | | | | | 12µm vs. 24µm | | ns | >0.9999 |
| | | | | | 12µm vs. Smooth | | ns | 0.1318 |
| | | | | | 24µm vs. Smooth | | ns | 0.3852 |
| Channe of variable complexity – Talin KO | 4d | 8 | na | 79 | 6µm vs. 12µm | One-way ANOVA with Kruskal-Wallis test with Dunn's post-test | ns | >0.9999 |
| | | | | | 6µm vs. 24µm | | ** | 0.0013 |
| | | | | | 6µm vs. Smooth | | **** | <0.0001 |
| | | | | | 12µm vs. 24µm | | * | 0.0188 |
| | | | | | 12µm vs. Smooth | | **** | <0.0001 |
| | | | | | 24µm vs. Smooth | | ** | 0.001 |
| Actin Flow in Channel of variable complexity – TIRF | 4g | 4 | na | 7 | 6µm vs. 12µm | One-way ANOVA with Friedman test with Dunn's post-test | ns | >0.9999 |
| | | | | | 6µm vs. 24µm | | * | 0.0423 |
| | | | | | 6µm vs. Smooth | | ** | 0.0036 |
| | | | | | 12µm vs. 24µm | | ns | 0.8499 |
| | | | | | 12µm vs. Smooth | | ns | 0.1649 |
| | | | | | 24µm vs. Smooth | | ns | >0.9999 |
| Channel Serrated – Control | Suppl.2c | 3 | 57 | na | 5-µm vs. 6-µm | One-way ANOVA with Kruskal-Wallis test with Dunn's post-test | ns | 0.7260 |
| | | | | | 5-µm vs. 7-µm | | ns | 0.2618 |
| | | | | | 5-µm vs. Smooth | | ns | 0.9653 |
| | | | | | 6-µm vs. 7-µm | | ns | 0.8396 |
| | | | | | 6-µm vs. Smooth | | ns | 0.4426 |
| | | | | | 7-µm vs. Smooth | | ns | 0.1117 |
| Channel Serrated – Talin KO | Suppl.2c | 4 | na | 9 | 5-µm vs. 6-µm | One-way ANOVA with Kruskal-Wallis test with Dunn's post-test | ns | >0.9999 |
| | | | | | 5-µm vs. 7-µm | | ns | 0.6027 |
| | | | | | 5-µm vs. Smooth | | ns | 0.0518 |
| | | | | | 6-µm vs. 7-µm | | ns | 0.5654 |
| | | | | | 6-µm vs. Smooth | | * | 0.0356 |
| | | | | | 7-µm vs. Smooth | | ns | 0.4275 |
| Actin Flow in confinement (EDTA) | Suppl.2f-g | 1 | 12 | na | TIRF actin flow | Column statistics: mean=13.72µm; Std Deviation=1.358µm; median=13.8µm | | |
| | | | | | TIRF+lamellipodia actin flow | Column statistics: mean=17.73µm; Std Deviation=1.265µm; median=17.88µm | | |
| | | | | | Cell lenght | Column statistics: mean=26.71µm; Std Deviation=2.674µm; median=27µm | | |

**Extended Data Figure 1**

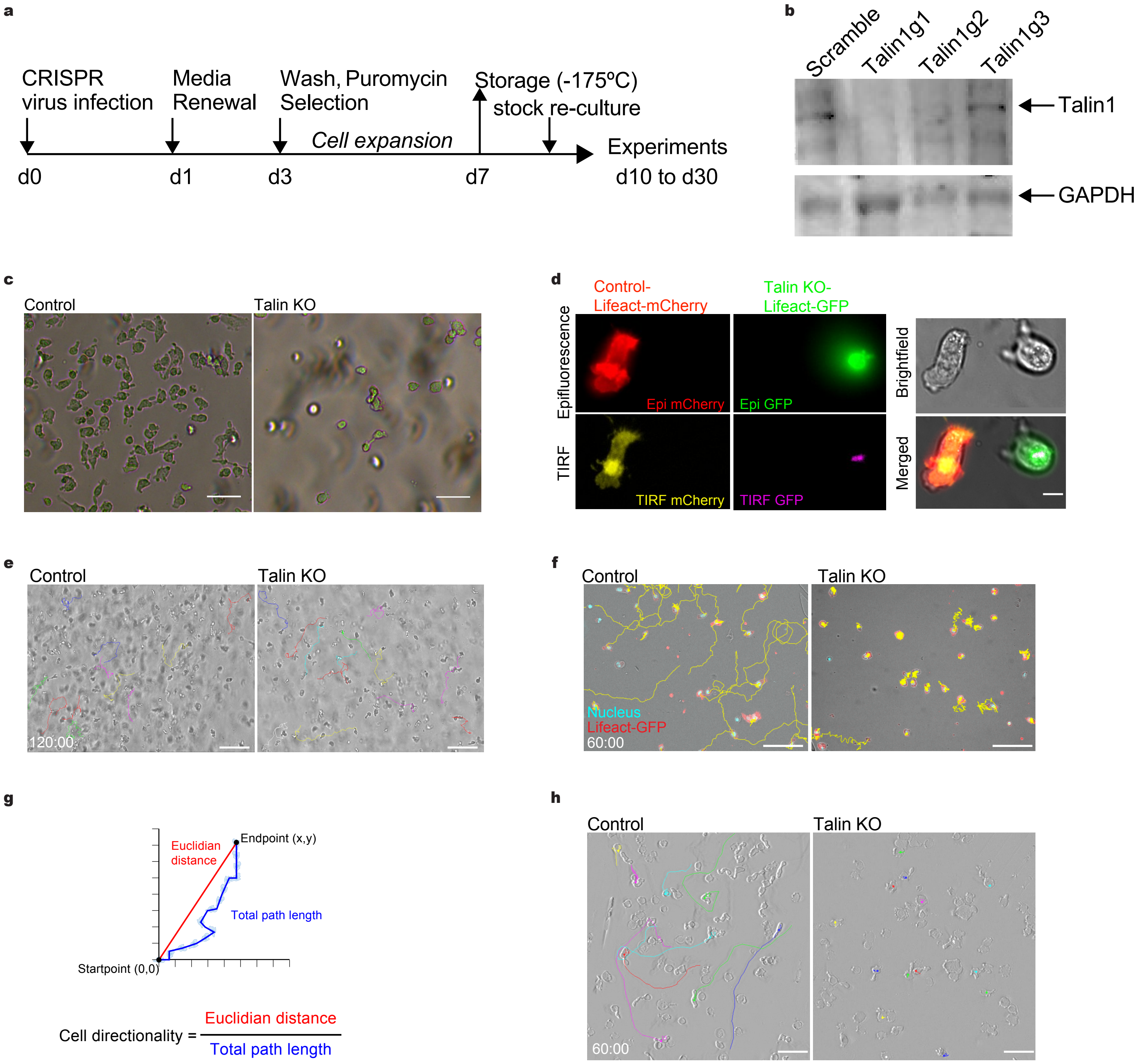

**Extended Data Figure 1 |**
**a**, Workflow scheme of virus production, cell infection, and analysis. **b**, Western blot analysis of T cells infected with lentivirus containing CRISPR guides scramble and talin1 (guide 1, 2 and 3). **c**, Phase contrast images of control and talin KO T cells focused on the 2D bottom of a tissue culture dish. Scale bars, 50 µm. **d**, Snapshots of epi-fluorescence and TIRF microscopy of control (left panel) and talin KO cells (middle panel) expressing either mCherry and eGFP reporters constructs for Lifeact, respectively, plated onto CD3/CD28-coated dish. Right panel shows bright-field and merged fluorescence, scale bar, 5 µm. **e**, Snapshots of control and talin KO T cell embedded in 3D collagen gel at t=60 min, individual tracks are displayed in colors. Scale bars, 100 µm. **f**, Control (left panel) and talin KO (right panel) T cells expressing Lifeact-eGFP (red), stained with Hoechst (cyan) were placed under 5 µm height confinement and imaged by video microscopy. Snapshot at t=60 minutes, nucleus tracks are display in yellow. Scale bars, 50 µm. **g**, Directionality is measured as the ratio of the displacement of the cell (Euclidian distance) by its total displacement length, to reduce the effect of the nucleus shifting within the cell on cell speed (Fig. 1d). **h**, Snapshots of control (left panel) and talin KO cells (right panel) at t=60 min. Individual tracks are displayed in colours, scale bars, 50 µm.

**Extended Data Figure 2**

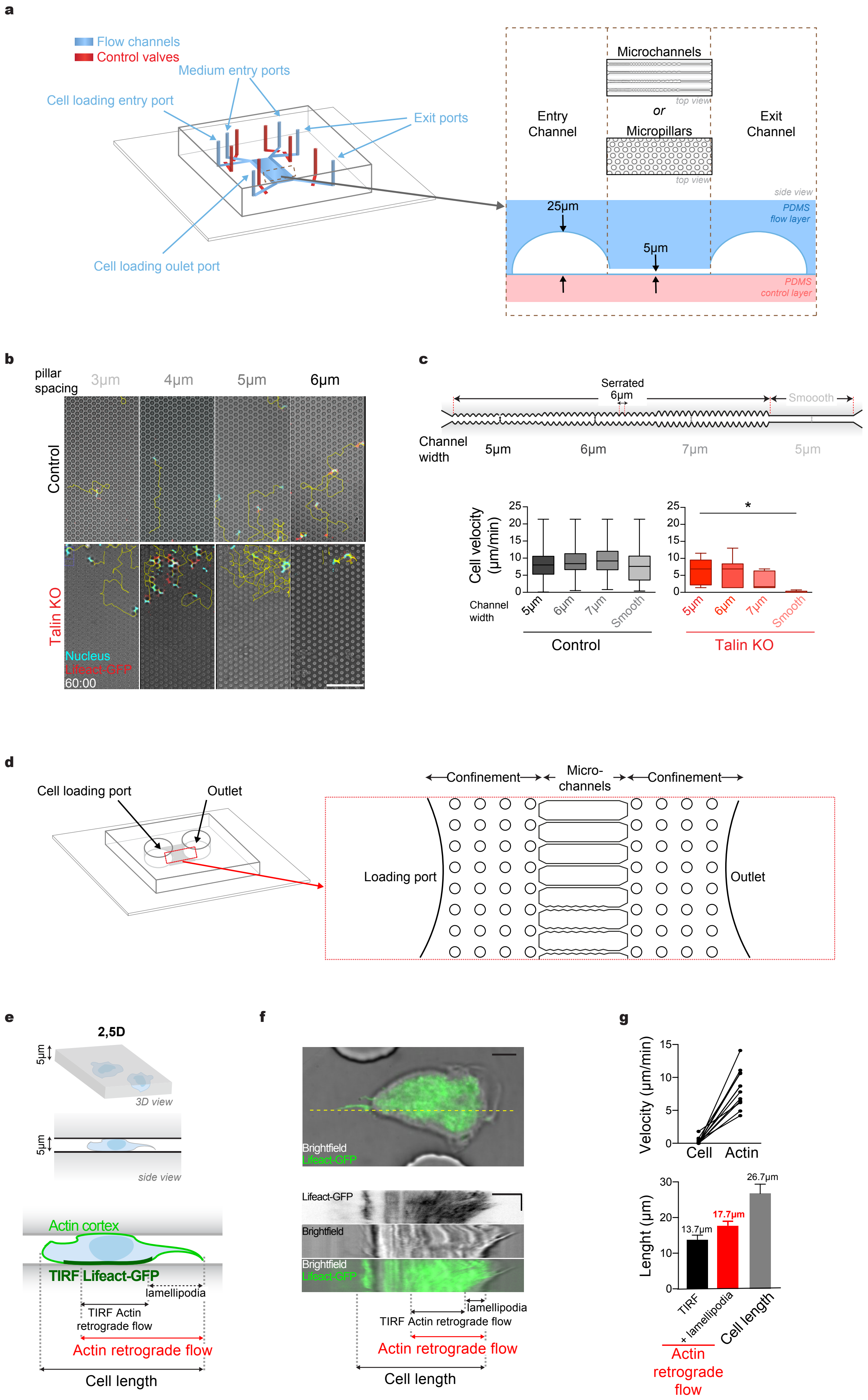

**Extended Data Figure 2 |**

**a**, Schematics of the multi-layered microfluidic devices used to push the cells into the pillar maze or into microchannels. **b**, Related to Fig. 2c-e: snapshots of a videomicroscopy at t=60 min of control and talin KO T cells in the maze device. Cyan shows the nucleus (Hoechst) and red the Lifeact-eGFP reporter, gray the brightfield image, and individual cell tracks are display in yellow. Scale bar, 50 µm. Time in min:sec. **c**, Related to Fig. 2i : cell velocities (µm/min) of control and talin KO T cell in serrated microchannels, where cell velocity in the serration zones of different diameters (with conserved spacing of 6 µm) are displayed (* p=0.0356, otherwise not significant; Kruskal–Wallis followed by a post hoc Dunn test). **d**, Scheme of the device used for the EDTA experiment. **e**, Measurement schematics of cell and actin retrograde flow length. Control T cells expressing Lifeact-eGFP reporter under 5 µm confinement with 10mM EDTA were observed by TIRF. **f**, Upper panel: snapshot of brightfield and TIRF. Bottom panel: kymographs along back-front polarity (yellow dotted line) of the cell. Arrows indicate the length of the cell, the retrograde actin flow observed by TIRF and the lamellipodia. Horizontal scale bars, 5 µm, vertical 1min. **g**, Upper panel: measurement of cell and actin retrograde flow velocities at a single-cell level. Lower panel: quantification of the mean cell length (26,7µm) and the actin retrograde flow length with (17,7µm) or without (13,7µm) the lamellipodium. Bars show standard deviation (n=12), one experiment.

**Captions for supplementary videos:**

**Supplementary video 1**: *Related to Fig. 1c.* Control (upper panel) and talin KO cells (lower panel) migrating in 3D collagen. Individual cell tracks are marked in colors. Time-lapse over 120 min, as indicated by the time stamp on the bottom left corner (h:min). Scale bar 100 µm.

**Supplementary video 2**: *Related to Fig. 1d.* Control (upper panel) and talin KO cells (lower panel) under 5 µm confinement. Cells are expressing Lifeact-eGFP reporter, displayed in red, and cell nucleus is marked in cyan with Hoechst 33342. Individual cell tracks are marked in yellow. Time-lapse of 60 min, as indicated by the time stamp on the bottom left corner (min:sec). Scale bar 50 µm.

**Supplementary video 3**: *Related to Fig. 1e-g.* Example of Total internal reflection (TIRF) of Control and talin KO T cells expressing Lifeact-eGFP under 5 µm confinement. First two examples display brightfield (left panel), Lifeact-eGFP TIRF in black (middle panel) and merged images (brightfield in gray, TIRF in green, right panel) acquired every 3 seconds. Example 3 present the cells shown in *Fig. 1E-G*, where TIRF of Lifeact-eGFP (black) was acquired every second. Time stamps in min:sec. Scale bars 5 µm.

**Supplementary video 4**: *Related to Fig. 2f-g.* Control (left panel) and talin KO T cells (right panel) migrating under 3 µm confinement. Individual cell tracks are highlighted with colors. Time-lapse of 60 min, as indicated by the time stamp on the bottom left corner (h:min). Scale bar 50 µm.

**Supplementary video 5**: *Related to Fig. 2a-e.* Control and talin KO T cells (upper and lower panel, respectively) migrating under 5 µm confinement bearing pillar arrays with separation of 3, 4, 5 and 6 µm. Cells are expressing Lifeact-eGFP reporter, display in red, and cell nucleus is marked in cyan with Hoechst 33342. Individual cell tracks are marked in yellow. Time-lapse of 60 min, as indicated by the time stamp on the bottom left corner (min:sec). Scale bar 50 µm.

**Supplementary video 6**: *Related to Fig. 2h-i.* Control and talin KO T cells (upper and lower panel, respectively) in 5x5 μm smooth microchannel. Cells are expressing Lifeact-eGFP reporter, display in red, and the cell nucleus is marked in cyan (Hoechst 33342). Individual cell tracks are marked in yellow. Time stamp on the bottom left corner in min:sec. Scale bar 20 μm.

**Supplementary video 7**: *Related to Fig. 2j-k.* talin KO and control T cell in 5x5 μm serrated microchannel. Cells are expressing Lifeact-eGFP reporter, displayed in red, and the cell nucleus is marked in cyan (Hoechst 33342). Note how the control cells manage in the smooth zone whereas the talin KO cells are blocked. Time-lapse of 60 seconds, as indicated by the time stamp on the bottom left corner (min:sec). Scale bar 20 μm.

**Supplementary video 8**: *Related to Fig. 2l-m.* Control T cells migrating in various microchannels, before and after addition of 10 mM EDTA. Cells are expressing Lifeact-eGFP reporter, displayed in red, and the cell nucleus is marked in cyan (Hoechst 33342). Individual cell tracks are marked in a speed-dependent color code (red: fast; blue:slow). Time-laps of 60 seconds, time stamp in min:sec. Scale bar 20 μm. 10 mM EDTA was added after imaging 90 min. Note that T cells can migrate in all conditions: confinement, serrated and smooth channel, and EDTA addition in the medium allow only fast cell migration in the serrated channels.

**Supplementary video 9**: *Related to Fig. 4a-d.* Control and talin KO T cell in 5 μm-high serrated microchannel with different periods between the serrations (6 μm, 12 μm, 24 μm and no serration). Cells are expressing Lifeact-eGFP reporter, displayed in red, and the cell nucleus is marked in cyan (Hoechst 33342). Individual cell tracks are marked in a speed-dependent color code (red: fast; blue:slow). Note how the talin KO cell can migrate in the highly serrated zones (6 and 12 μm distance) and not in the low serration part (24 μm distance and no serration). Time-lapse of 60 seconds indicated by the time stamp (min:sec) on the lower left corner. Scale bar 20 μm.

**Supplementary video 10**: *Related to Fig. 4e-g.* Talin KO T cell expressing Lifeact-eGFP reporter in 5 µm-high serrated microchannel with different periods between the serrations (6 µm, 12 µm, 24 µm and no serration) with high magnification (63X), in merged brightfield and TIRF (upper panel, gray and green, respectively) and TIRF (lower panel, in black). Time-lapse of 2 seconds indicated by the time stamp (min:sec) on the lower left corner. Note how the cell migrates in the highly serrated zones (6 and 12 µm distance) and not as efficiently in the low serration part (24 µm distance and smooth), where it displays a higher actin retrograde flow. Scale bar 10 µm.
